# Digital Cell Sorter (DCS): a cell type identification, anomaly detection, and Hopfield landscapes toolkit for single-cell transcriptomics

**DOI:** 10.1101/2020.07.17.208710

**Authors:** Sergii Domanskyi, Alex Hakansson, Thomas Bertus, Giovanni Paternostro, Carlo Piermarocchi

## Abstract

**Motivation:** Analysis of singe cell RNA sequencing (scRNA-seq) typically consists of different steps including quality control, batch correction, clustering, cell identification and characterization, and visualization. The amount of scRNA-seq data is growing extremely fast, and novel algorithmic approaches improving these steps are key to extract more biological information. Here, we introduce: (i) two methods for automatic cell type identification (i.e. without expert curator) based on a voting algorithm and a Hopfield classifier, (ii) a method for cell anomaly quantification based on isolation forest, and (iii) a tool for the visualization of cell phenotypic landscapes based on Hopfield energy-like functions. These new approaches are integrated in a software platform that includes many other state-of-the-art methodologies and provides a self-contained toolkit for scRNA-seq analysis.

**Results:** We present a suite of software elements for the analysis of scRNA-seq data. This Python-based open source software, Digital Cell Sorter (DCS), consists in an extensive toolkit of methods for scRNA-seq analysis. We illustrate the capability of the software using data from large datasets of peripheral blood mononuclear cells (PBMC), as well as plasma cells of bone marrow samples from healthy donors and multiple myeloma patients. We test the novel algorithms by evaluating their ability to deconvolve cell mixtures and detect small numbers of anomalous cells in PBMC data.

**Availability:** The DCS toolkit is available for download and installation through the Python Package Index (PyPI). The software can be deployed using the Python import function following installation. Source code is also available for download on Zenodo: doi.org/10.5281/zenodo.2533377

## 1 Introduction

Several platforms have emerged over the last decade for high-throughput RNA sequencing at the single-cell resolution (32). Obtaining biological insights from single-cell data, however, requires complex computational analysis. Generally, the first part of the analysis includes quality control (QC) of raw base call (BCL) files and alignment of reads with the reference genome, followed by their quantification at the gene-level. The second part of the analysis starts with additional QC procedures to filter out cells with low library size and low number of unique genes sequenced, detect multiples (i.e two or more cells are captured instead of one), and remove cells with high content of mitochondrial genes, which indicates broken cells (16). Data is then normalized to ensure that the gene expression levels are comparable between individual cells. Existing normalization methods (9) have different strengths and weaknesses. It is therefore important that scRNA-seq software provides the flexibility to choose the best normalization strategy for a given dataset. After normalization, the most variable genes’ largest principal components from Principal Component Analysis (PCA) are often used for clustering and two-dimensional visualization. After clustering, biological insight is obtained using different methods, including differential expression, network analysis, enrichment analysis and cell type annotation.

For users, it is most desirable to have an extensive toolkit in one software package. Existing packages for scRNA-seq data include Cell ranger (31), edgeR (23), DESeg2 (20), Seurat (25), Scanpy (30), but many others are available. The advantages of an extensive set of methods under one open-source software umbrella are smooth integration of the processing steps and control of the workflow. For example, Seurat is a powerful package implemented in R with large data integration capability and visualization, Scanpy is a Python-based software specifically designed for efficient processing of large single-cell transcriptomics datasets. In addition, commercial providers of single-cell platforms have developed software tools for low-level processing and visualization (11), focusing on the first part of the analysis, i.e. up to generating the gene counts matrix.

Here, we join the ongoing effort to provide the community with user-friendly tools for the second part of single-cell transcriptomics analysis, starting with pre-processing and quality control and ending with cell type annotation, visualization, and analysis of the annotated clusters. In addition to state-of-the-art methods, our *Digital Cell Sorter* (DCS) platform introduces three new algorithms: (1) An enhanced version of our recently-developed algorithm for the automatic annotation of cell types, *polled DCS* (p-DCS) (10), which uses a predefined set of markers to calculate a voting score and annotate cell clusters with cell type information and its statistical significance. The enhanced version of p-DCS uses a marker-cell type matrix normalization method to account for markers that are known to be unexpressed in certain cell types. Also, in the new version, low quality scores are set to zero to reduce noise in cell type assignment. (2) A tool that provides a cell anomaly score using isolation forest, an unsupervised algorithm ((18)) for anomaly detection. While clustering is based on similarity, our cell anomaly score detects and quantifies the degree of heterogeneity within each cluster and is important, for instance, in the analysis of scRNA-seq from cancer samples. We also use this score to detect cells that are different than the majority of the other cells in a dataset, like in the case of anomalous circulating cells in blood. (3) A second algorithm for cell type identification based on Hopfield networks ((15)). Hopfield networks allows for a direct mapping of associative memory patterns, in this case patterns of gene expression, into dynamic attractor states of a recurrent neural network. This method has been successfully used in the classification of cancer subtypes ((6, 8, 21, 26, 29)). In our algorithm, we use cell type markers to define Hopfield attractors, and we let clusters of cells evolve to align with these attractors. The Hopfield network is integrated with an underlying biological genegene network, the Parsimonious Gene Correlation Network (PCN) ((7)), to retain only biologically significant edges. This allows us to obtain interpretable information on the role of specific markers and their local connectivity in defining the different cell types. The method also defines an energy-like function that permits the visualization of the gene expression landscape and represents cell types as valleys associated to the different cell type attractors.

The different tools in the DCS platform can be combined for improved performance. For instance, we show how to combine the methods in (1) (p-DCS) and (2) (Hopfield classifier) into a consensus annotation methodology that is more accurate compared to the methods used separately. Finally, we provide examples of the capabilities of DCS and its performance using data from large single cell transcriptomics datasets of peripheral blood mononuclear cells, and bone marrow samples from healthy and multiple myeloma patients. Note that our cell annotation methods are knowledge-based classifiers, since they rely on pre-existing knowledge from cell type markers and do not require training data.

## 2 Methods

### 2.1 Functionality overview and toolkit structure

DCS functionalities include: (i) pre-processing (handling of missing values, removing all-zero genes and cells, converting gene index to a desired convention, normalization, log-transforming); (ii) quality control and batch effects correction; (iii) cells anomaly score quantification; (iv) dimensionality reduction and clustering; (v) cell type annotation; (vi) visualization, and (vii) post-processing analysis.

We classify our tools into three categories: primary processing tools, data query tools, and visualization tools. Primary processing tools consist of functions transforming input files into valid tables and translating gene names to a desired convention, general data pre-processing, dimensionality reduction, and clustering. Cell type identification and anomaly score calculations are also included in the processing tools. Data query API (Application Programming Interface) tools include functions for retrieving gene expression of one or many genes across all cells or in a subset of cells, extraction of new marker genes characteristic of a cell cluster, and other querytype functionalities. Visualization tools include two-dimensional projection, quality control histograms, marker expression projections, marker expression summaries, gene expression heatmaps, individual gene t-texts, cell types assignment matrices, cell types stacked barplots, anomaly scores projections, pDCS null distribution histograms, new markers plots, Sankey diagrams (a.k.a. river plot), and cell type markers summary diagrams.

### 2.2 Cell type annotation

After unsupervised clustering, DCS automatically assigns clusters to cell types without relying on an expert curator to interpret the data. DCS uses all the information available in a knowledgebase of characteristic markers for many cell types. While cell type identification by manual interpretation generally provides good results, DCS assures that all the available information, including the presence and absence of markers, is taken into account, and can automatically identify cell types in very large datasets. Moreover, DCS provides statistical significance of the assignments, labeling as unknown clusters for which there is not sufficient information to make an assignment, and provides multiple labels with their associated scores when the expression pattern is consistent with more than one cell type. In DCS, automatic cell type annotation can be obtained using two methods: a voting algorithm and an Hopfield recurrent network classifier.

#### 2.2.1 Voting algorithm

The voting algorithm in DCS is based on an extensive revision of our polled Digital Cell Sorter method ((10)). Prior information on cell markers is encoded in a marker/cell type matrix *M_km_* where *k* is the cell type, and *m* is the marker gene. The element *M_km_ =* ±1 if *m* is an expressed/not expressed marker of cell type *k.* We will refer to these as positive and negative markers, respectively. Finally, if marker *m* is not used in determining cell type *k* the corresponding element of *M* is zero. We normalize *M*, separately for negative and positive markers, by the number of markers expressed in each cell type and then by the number of cell types expressing each marker. Thus markers that are unique to a particular cell type will be automatically assigned a large weight. By retaining markers in *M* that are expressed in a given dataset *X*, we obtain a matrix 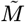. In Fig. 1 (bottom panel) we show an example of 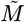 for a PBMC dataset ((31)), after gene filtering and normalization, with dark green (rose) corresponding to unique positive (negative) markers.

**Fig. 1.**
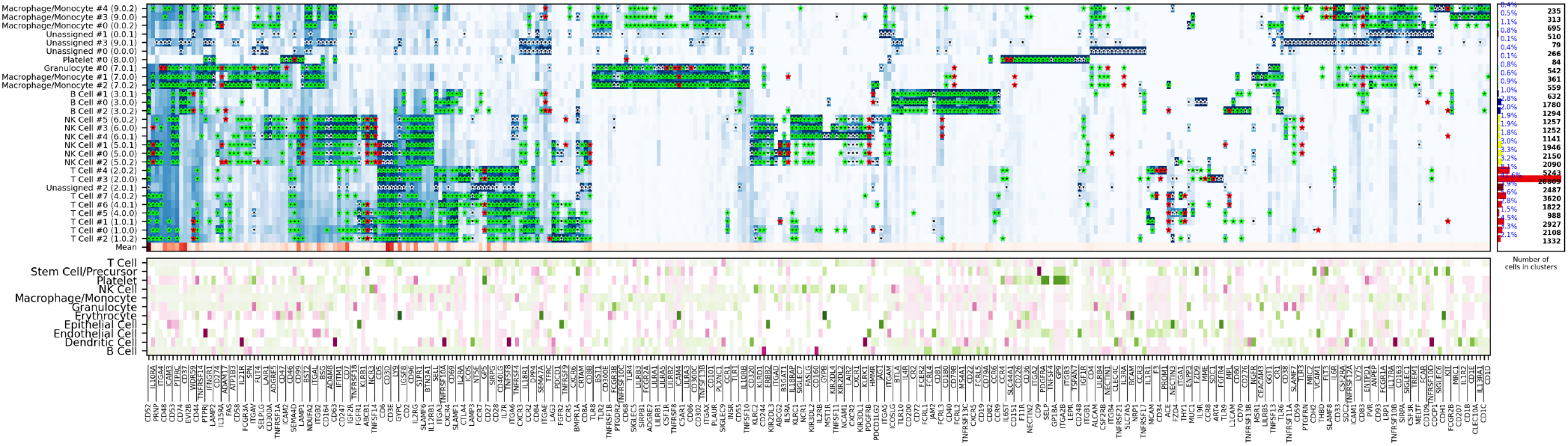
Top panel: heatmap of the average gene expression for each cluster, with darker blue corresponding to higher expression values. Green stars indicate supporting markers of the assigned cell type, red stars are contradicting, and white stars are neither supporting nor contradicting but significantly expressed. The assigned cell type is indicated on the left and combined with the cluster number id. Bottom panel: normalized marker cell type matrix *M* with dark green (rose) indicating unique positive (negative) markers. Right panel:total numbers of cell for each type and cluster.

We then build the marker/centroid matrix *Y_mc_* of the mean expression of marker *m* across all cells in cluster *c.* For each marker *m*, we use *Y_mc_* to compute all cluster centroids’ z-scores *Z_mc_.* The z-score matrix *Z_mc_* is transformed into the matrix 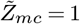 if 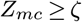 and 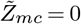 otherwise, for a given threshold *ζ*. The number of possible supporting markers decreases by increasing the value of the cutoff *ζ*, and this parameter has to be selected so that each cell type expected to be present in the dataset has a sufficient number of markers. Fig. 1 (top panel) shows *Y_mc_,* calculated for the PBMC dataset, with darker blue corresponding to higher expression of markers, and stars denoting statistically significant markers, i.e. markers with z-score larger than *ζ*. We have varied the parameter *ζ* in the range 0.1-1.5, and for the dataset in the figure, we chose *ζ* = 0.3. Finally, we compute the matrix of voting scores for each type-cluster pair (*k*, *c*) according to 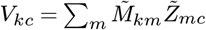.

To quantify the statistical significance of the voting scores and make the final assignment, we use a randomization method to calculate the statistical uncertainty associated to each type-cluster pair (*k*, *c*). We randomize the clusters by preserving their size and assigning to them cells randomly chosen from the whole dataset, and compute the voting scores for each random configuration. This randomization is performed *n* = 10^4^ times, recording the voting matrix *V_kc_* for each configuration of random clusters. This method accounts for cluster sizes, the overall gene expression distribution of the markers, and imbalances in the number of markers per cell type in estimating the uncertainty. The procedure provides distributions of voting results 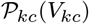 for a null model of random clusters.

We determine the z-scores, Λ*k_c_*, of the voting results *V_kc_* in the null distribution 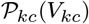 and assign the cell type according to *T_c_ =* argmax_*k*_ Λ_*kc*_. All cells belonging to cluster *c* are thus identified as cell type *T_c_*. In the example of Fig. 1, we show in the top panel how, after the cell type of each cluster has been assigned, we use green/red stars to indicate supporting/contradicting markers. The labels assigned to clusters are indicated on the left side of the top panel, and the total numbers of cell of each type and cluster are shown in the right panel.

DCS also includes an algorithm that consists in a modification of the above voting method, and is similar to a recently-proposed evidence-based cell-type identification algorithm (24). In this modification, we divide the voting scores *V_kc_* by the maximum possible scores that each cell type could have if all positive makers and none of the negative markers were expressed,

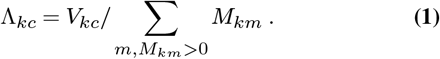

We then assign the cell type according to *T_c_* = argmax_*k*_ Λ_*c*_. The main difference with respect to the algorithm in (24) is that we account for negative marker genes as well as marker weights. Both annotation methods introduced in this section can be combined to obtain a consensus score using the geometric mean of the corresponding Λ_*kc*_.

We remark that, in contrast to the previous version of our algorithm ((10)), we now account for negative markers, i.e. markers that should not be significantly expressed, in the cell type assignment. The normalization procedure has been modified accordingly. A marker can be in one of three states: supporting, contradicting, or neither supporting nor contradicting. Moreover, to discard low quality score, we now set the score to zero if the number of supporting markers is below 10% of all known markers for a given cell type.

#### 2.2.2 Hopfield knowledge-based classifier

Cell type assignment with this algorithm is based on the idea that gene expression for different cell types can be represented as *associative memories,* and encoded as attractor states of the signaling dynamics in an underlying gene network. If one starts with cells in a cluster and let them evolve according to the dynamics defined by a set of interacting memory patterns, the overlap of the cluster configuration with the attractors can be used for cell type assignment. Hopfield networks ((15)) are the simplest models of associative memories and are defined using *N* Boolean variables *σ_i_* (*t*) evolving at integer time steps *t*. In our case these variables are associated with the expression of each gene. The initial state of each node (gene) takes one of two values, *σ_i_* (*t*) = ±1 (over/underexpressed), based on the statistical significance of the average marker expression in each cluster, determined as a z-score above a threshold *ζ* = 0.3 across all clusters. In the canonical Hopfield model, a coupling matrix is constructed to store a set of *p* independent Boolean patterns 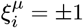 as point attractors, where *i* = 0,1,…,*N* – 1 is the node index and *μ* = 0,1,…,*p* – 1 is the pattern index. In the algorithm implemented in DCS, we build attractors 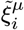 using our normalized marker cell type matrix, 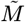, detailed in Sec. 2.2.1. The negative elements of this matrix are set to zero, thus negative markers are not used in this method. The coupling matrix *J_ij_* defines the strength and sign of the signal sent from node *j* to node *i* and is defined by

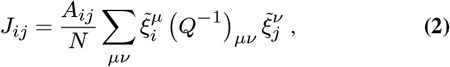

where

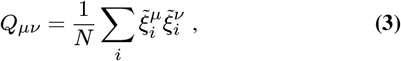

is a matrix that reduces the effects of correlation in the attractors ((2, 3)), and *A_ij_* is the adjacency matrix of the underlying biological gene-gene interaction network. DCS currently uses a generic gene correlation newtwork, the Parsimonious Gene Correlation Network (PCN) ((7)), which can be easily replaced with other gene networks. The underlying network defined by *A_ij_* effectively reduces interactions between the nodes of the Hopfield neural network to retain only biologically-justified gene-gene interactions.

The total field at node *i* at time *t* is given by

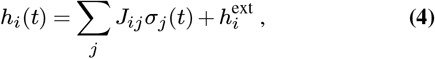

and the dynamical update rule is given by

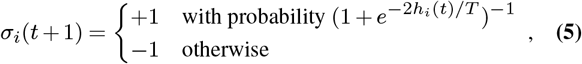

where *T* is an effective temperature representing noise (not a physical temperature). Biologically, this noise represents the effect of different kinds of biochemical fluctuations in cells. Additionally, any node in the network that is being turned off, i.e. the local field on a given node is such that it would switch from +1 to −1, is instead set to 0. This simple modification to the dynamical rule above makes the signaling dynamics dependent not only on the neighboring genes, but also on the gene state at the previous step, and the network becomes more diluted during the evolution. Moreover, this way of updating the system is asymmetric since it affects only genes that switch from +1 to −1, and not, for instance, nodes that stay from −1 to −1. This modified Hopfield dynamics has been recently proposed by (6), and we have found, like they did, that this rule greatly improves the convergence of the classifier. This modification is useful because nodes correctly overexpressed in the input and associated to a +1 get sometimes switched to −1 only due to the stochastic nature of the algorithm. Removing these nodes avoids the amplification of these wrong switches in the following updating steps.

The update rule from Eq. 5 may be implemented in various ways. The synchronous scheme updates the state of all the nodes in the system at every time step, but this is sensible only if the simulated system has a central pacemaker coordinating the activity of all nodes. A more appropriate choice for decentralized systems is the asynchronous scheme, in which the state of a randomly chosen subset of nodes is updated at each time step. Here, we use the asynchronous scheme with update probability for each node that linearly increases from 0.025 to 0.5 in the course 100 time steps, after which it is kept constant.

The overlap of the state vector *σ_i_* (*t*) with the *μ*^th^ pattern in one of the attractor states is given by

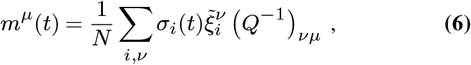

where −1 ≤ *m^μ^* (*t*) ≤ +1. The overlap measures the similarity between the gene expression of the different attractors and the simulated gene expression, and *m^μ^* (*t*) = +1 means that there is perfect agreement between the simulated expression and attractor pattern *μ.* For each cluster we assign the cell type corresponding to the maximum non-negative overlap with the attractor states at the last time point. If for a given cluster there are no non-negative overlaps with any of the attractor states, then a “Unassigned” label is assigned. The stochastic nature of evolution of Hopfield network may lead to slightly different dynamics between independent realization, therefore the simulation is repeated multiple times and the most frequently assigned cell type is selected as a label for each cluster. Second, third, and other most frequent cells types are also recorded into the score matrix with their corresponding frequencies.

The two methods for cell assignment in the previous section and the Hopfield classifier can be combined in different ways to obtain a consensus approach using the geometric averages of the corresponding score matrices. All combination options are detailed in the DCS documentation.

### 2.3 Hopfield landscapes visualization

The Hopfield model introduced in the previous section can be used to define a quasi-energy function

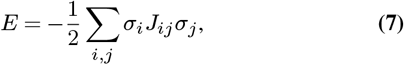

where *σ_i_* is the Boolean variable describing gene up or down regulation, and the matrix *J* is defined using Eq. 2. Eq. 7 defines a Lyapunov function for the signaling dynamics in the gene network and allows us to build a *Hopfield landscape* in which the attractor states, i.e. the different cell types, are the minima of a complex multi-dimensional phenotypic landscape. The quasi-energy in Eq. 7 can be defined with or without the matrices *A_ij_* and *Q_μv_*, and using different normalizations for the 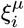. Therefore, our DCS software allows for different options for the landscape representation. This landscape can be explicitly visualized by starting at the equilibrium points corresponding to the attractor configurations and adding noise to sample their basins of attraction. The points sampled can then be represented in a 2D plot using principal components projections ((12, 21, 27, 28)). An example of output from this DCS visualization tool is in Fig. 2, where the *A_ij_* was used (and not the *Q_μν_*) and the 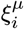 were normalized as in the table 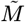. (See Supplementary Fig. 1 for a landscape with *Q_μν_*). The figure shows the visualization of the phenotypic landscape of 14 different hematopoietic cell types, where the coordinates are the first two largest principal components of the point attractors, and the colors and contour lines reflect the Hopfield quasi-energy function of Eq. 7. These attractors were built using markers from ((22)), modified to merge subtypes of B cells, CD4 T cells, NK cells, Macrophages, Dendritic cells and Mast cells, and their position in the landscape are indicated by stars. This visualization effectively uses the Hopfield quasi-energy to represent the matching/mismatching with the attractors in all dimensions. Note how for some cell types the basins of attraction are closer to each other than for others, and sometimes they overlap. For instance, Mast Cells, Eosinophils, and Plasma cells have overlapping basins, well separated from other cell types such as T cells or Macrophages. Additional considerations and analyses on Hopfield attractors and the role of the *Q* matrix are discussed in Supplementary Materials (Supplementary Figs.1-2).

**Fig. 2.**
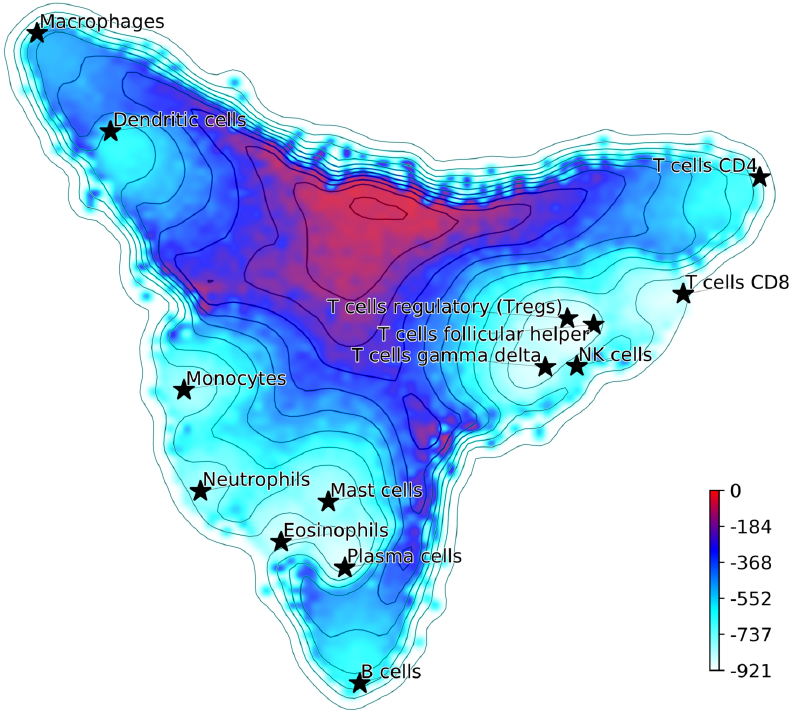
Hopfield attractor landscape visualization. The points are colored according to their Hopfield quasi-energy, Eq. 7.

### 2.4 Cell anomaly quantification

This module implements an algorithm that quantifies cell anomaly, i.e. how much cells are different from other cells within the same dataset or within one or more clusters. Anomaly detection and can be interpreted as the opposite of clustering, which is based on similarity measures. The module for the quantification of anomaly is based on the Isolation Forest anomaly detection algorithm((18, 19)).

The Isolation Forest algorithm isolates cells by randomly selecting a gene and a split value for its expression value. A schematic illustration of the algorithm is shown in Fig. 3. For the sake of illustration, consider a genome with two genes A and B only (along the two axes in Fig. 3 (a)). Random partitioning is illustrated by the vertical and horizontal lines labelled with *s*_1_,…,9. At each step the cells are partitioned in two sets by choosing one of the two genes and a random threshold for the chosen gene’s expression. For a typical (green) cell (Fig. 3 (a)), the number of steps necessary to isolate the cell is larger than for an anomalous cell (red). This recursive partitioning can be described by a tree, as shown in Fig. 3 (b), where each internal node contains a gene and a threshold value that were used in that partitioning, whereas leaf nodes are the isolated cells. The partitioning is carried out until all cells are isolated, i.e. no further partition is possible. The cell anomaly algorithm has a training and an evaluation stage. In the training stage, an ensemble of *n* = 100 isolation trees, called isolation forest, is generated using random sub-samples of *ψ* = 256 cells from the original dataset. Next, in the evaluation stage, for each cell *i* in the dataset, or in a subset of cells of interest, and for each tree *j* in the forest, a path length *h_ij_* is derived by counting the number of edges from the root node of the tree to the node where the cell is isolated by following genes and thresholds stored in tree *j*. To compute anomaly score of cell *i*, path lengths *h_ij_* obtained from all *n* trees in the isolation forest are averaged and normalized according to

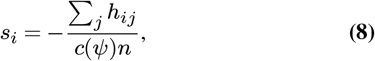

where *c*(*ψ*) is a normalization factor that accounts for the subsampling size *ψ*, giving a single score for the cell *i*, or a measure of anomaly. The algorithm’s pseudocode, and details about normalization factor, special conditions in the anomaly score computation and optimal selection of *n* and *ψ* are discussed in the original publication of the anomaly detection algorithm ((18)). When a forest of random trees collectively results in shorter branches for particular cells, they are likely anomalous cells. Cell anomaly can be used to rank cells from more anomalous to less anomalous. Then the top-ranked cells can be further analysed to investigate the biological difference from the other cells in the dataset. This module can be useful to identify small numbers of cells that may not be otherwise separated into a distinct cluster.

**Fig. 3.**
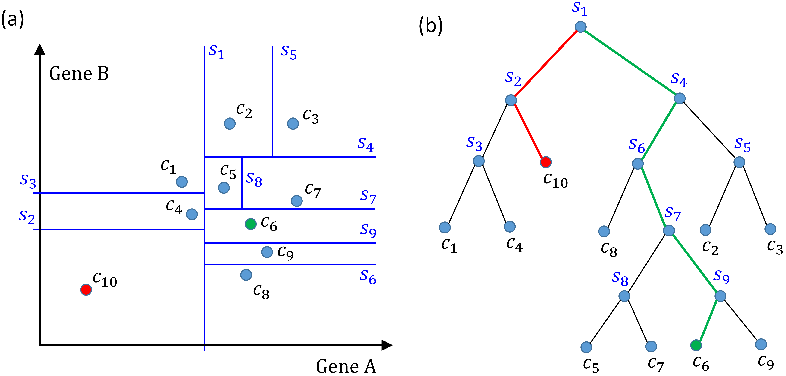
Schematics of how the isolation forest algorithm quantifies cell anomaly. In this example we consider only two genes A and B. One of these two genes is randomly selected and data is partitioned based on a random threshold applied to the chosen gene’s expression (vertical and horizontal lines labelled with *s_i_*,…,9). Common cells, e.g. *c*_θ_ in (a), will be isolated with a larger number of partitions compared to anomalous cells, e.g. *c*_10_. (b) Tree representation of the algorithm, where each internal node contains a gene and a threshold and represents a partition of the cells. Random partitioning produces shorter branches for anomalous cells (red) compared to common cells (green).

### 2.5 Quality control

Supplementary Fig. 3 shows an example of output from the quality control module. Count depth, i.e. number of reads per gene, and gene counts, i.e. number of non-zero genes per cell, are evaluated for each cell. Cell with count depths and gene counts that are below 50% of the median of their respective distributions are tagged as low quality. The quality control parameters defaults can be overwritten by a software user, as different datasets may require adjustment of the cutoffs.

The DCS quality control procedure also accounts for the fraction of mitochondrial genes expressed in each cell, as shown in Supplementary Fig. 3 (c). The fraction of mitochondrial genes cutoff is chosen at the abscissa interception with a line that passes through the median of the distribution and median plus 1.5 standard deviations. The list of human mitochondrial genes is taken directly from MitoCarta2.0: an updated inventory of mammalian mitochondrial proteins ((5)).

The integration of DCS with other state-of-the-art algorithms for batch correction, clustering, dimensionality reduction and other visualization tools is detailed in the Supplementary Materials (Supplementary Figs. 4-7).

## 3 Results

### 3.1 Cell identification in mixtures

We compared the performance of our previous p-DCS method ((10)), with the new Hopfield classifier and the consensus annotation classifier described in Section 2.2. To evaluate the algorithms’ performance, we randomly generated 100 mixtures of pure cell types, representing a gold standard, and evaluated the algorithms based on their automatic annotation. As gold standard, we used data from CD14 monocytes, CD19 B cells, CD34 cells, CD4 memory T cells, CD4 naive T cells, CD4 regulatory T cells, CD4 helper T cells, CD8 cytotoxic T cells, CD8 naive cytotoxic T cells and CD56 NK cells from a FACS-sorted PBMC dataset ((31)), in addition to endothelial (SRS2397417, SRS4181127, SRS4181128, SRS4181129, SRS4181130) and epithelial cells (SRS2769050, SRS2769051) annotated in the PanglaoDB database ((13)). The total number of cells used was 98,752, 53% of which are T cells. Each independent synthetic mixture contains on average 5,000 cells, chosen randomly from the 16 cell types listed above in random fractions. As the algorithms uses prior knowledge in the form of marker genes, we tested the algorithms using two marker genes tables: (i) CD Marker Handbook containing 11 main cell types and covering all 16 cell-types and sub-types present in the gold standard set ((4)), and (ii) CIBERSORT LM22 ((22)), modified to merge subsets of B cells, T cells, NK cells, Macrophages, Dendritic cells and Granulocytes. Marker table (ii) does not have marker information for CD34 cells, Endothelial cells or Epithelial cells, which are 25.6% of all the cells. To quantify each method’s performance we calculated a multi-class weighted F1 score, excluding unassigned cells, and took the median value over 100 independent random mixtures. We also calculated the median of the fraction of cells that had cell type unassigned for the same random mixtures. The results are shown in Tab. 1. Note how, for both marker tables, the new Hopfield classifier performed better than the original pDCS method and that the consensus annotation outperformed all other methods. Recently, pDCS has been independently compared to other methods for automatic cell identifications by (1), showing that pDCS was the second best algorithm in the prior-knowledge category.

**Table 1.**
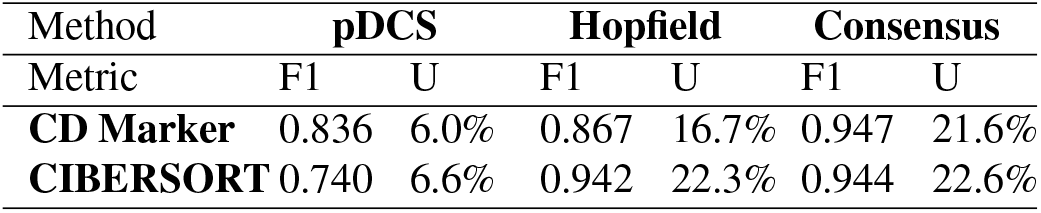
Performance of annotation methods evaluated using mixtures of pure cell type populations. Median multi-class F1 and percentage of unassigned cells (U) using two lists of markers as knowledge-base.

| Method | pDCS |  | Hopfield |  | Consensus |  |
| --- | --- | --- | --- | --- | --- | --- |
| Metric | F1 | U | F1 | U | F1 | U |
| **CD Marker** | 0.836 | 6.0% | 0.867 | 16.7% | 0.947 | 21.6% |
| **CIBERSORT** | 0.740 | 6.6% | 0.942 | 22.3% | 0.944 | 22.6% |

A particular case of cell identification in a mixture is shown in Fig. 4. This figure shows how the platform can be used to analyze complex mixtures involving different tissues. Here we combined scRNA-seq data from: (1) CD138+/CD38+ cells from bone marrow samples obtained from hip replacement surgery ((17)) and (2) PBMC ((31)). The samples from (1) are dominated by plasma cells due to the CD138+/CD38+ pre-selection by flow cytometry, while plasma cells should be relatively rare in the PBMC sample. We therefore expect our algorithm to label most of the cells from (1) as plasma cells, and identify other cell types in the cells coming from (2). Fig. 4 (a) shows the t-SNE layout of the mixture of 8448 CD138/+CD38+ cells and 1000 PBMCs randomly selected cells from the PBMC dataset, after QC and batch effect correction. Clusters annotated with cell type information are shown in Fig. 4 (b) and the relative size of the clusters are shown in panel (c), including cells that did not pass QC. Annotation was obtained using the CIBERSORT LM22 list as knowledge-base. Note how in this particular case the algorithm correctly identified 96% of the cells from (1) that passed QC as “Plasma cells”, while most of the remaining was assigned to “Monocytes”. Cells from (2) were mostly identified as T cells, which is the dominant cell type expected in dataset (2). The Sankey plot in panel (d) shows on the left side the clusters with their label and on the right side the batches from datasets (1) (labeled by “hip”) and (2). The thickness of the lines is proportional to the number of cells in the corresponding cluster-batch pair. Note how most of the cells from dataset (1) are linked to clusters labeled either as plasma cells or failed QC.

**Fig. 4.**
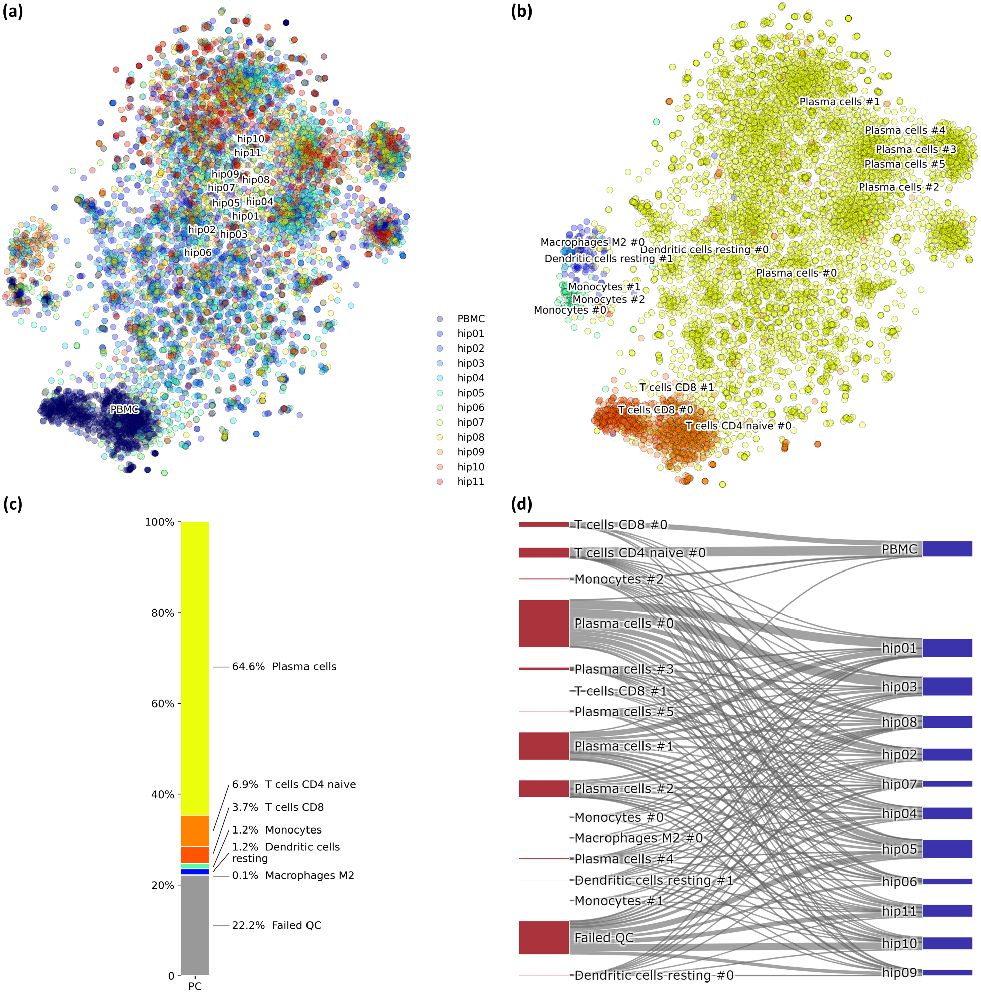
(a) Mixture of 8448 CD138+CD38+ cells and 1000 PBMCs on a t-SNE layout. Batch effects arising from different datasets, labelled by different colors, were removed with COMBAT. (b) Same layout as in (a) with clusters annotations. (c) Relative fractions of the annotated cell types, including cells that failed QC requirements. Such cells are excluded from plots in (a) and (b). (d) River plot showing labelled categories, from (b) and (c), and their memberships for each of the batches.

### 3.2 Performance with different marker lists

Since our voting and Hopfield-based algorithms depend on a list of marker genes, we have studied the dependence of cell type assignment on different lists of markers by comparing the labels assigned to a gold standard. In addition to the CIBERSORT and the CD Marker Handbook introduced above, we created a new list of markers obtained by differential expression analysis of data from a large recently-published single-cell study ((14)), and we also used a manually-curated list of cell-type markers from PanglaoDB ((13)). These cell marker lists differ in the number of markers per cell type, presence/absence of negative markers, and overlap of markers across cell types. More importantly, these lists are based on different labels, often corresponding to different degrees of type/subtype refinement. As a gold standard we used a set of 68579 PBMC cells from (31). Fig. 5 shows Sankey plots connecting cells annotated using the consensus annotation and one of the markers list (on the left side of each plot) with the annotation in the gold standard (right side of each plot). This visualization allows for a direct comparison of cells labeled using different degrees of type/sub-type refinement. The plot also explicitly indicates cells that were labeled as “Unassigned” and cells that did not pass QC. Overall, a comparison among these four plots indicates that the annotation is quite consistent.

**Fig. 5.**
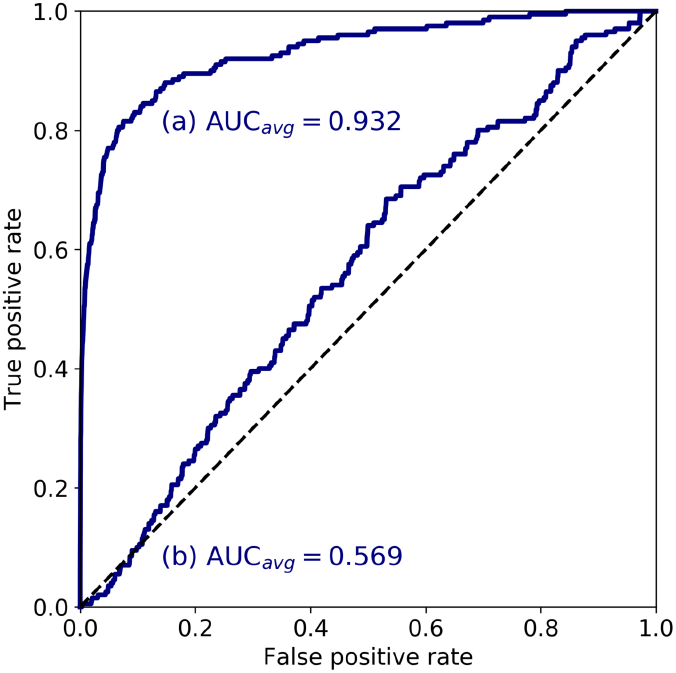
Sankey diagrams for 68579 PBMC annotated cells (taken as gold standard, (31)) using marker gene information from: (a) CIBERSORT LM22, (b) CD Marker Handbook, (c) HCL peripheral blood samples, (d) PanglaoDB. The plots connect cells annotated using our consensus annotation with one of the four marker lists (on the left side of each plot) with the annotation in the gold standard (right side of each plot).

### 3.3 Validation of anomaly detection

We have designed an *in silico* experiment to validate and demonstrate the utility of the cell anomaly score introduced in Sec. 2.4. The experiment mimics a scenario in which a small number of circulating endothelial cells are present in periferal blood, and evaluates the algorithm on its ability to detect these cells using their anomaly score. We selected 1637 cells (T cells and Monocytes of PBMC) from SRS3363004 (GSM3169075) and 70 endothelial cells from SRS3822686 (GSM3402081) to mimic the presence of cells that are rare in blood. Both datasets were annotated in PanglaoDB and have been sequenced using Illumina HiSeq 2500 and 10x Chromium. The two datasets have similar sequencing depth, median number of expressed genes per cell and fraction of good quality cells, as detailed in Supplementary Table 1.

We combined 1637 PBMC and 20 endothelial cells randomly chosen out of the total 70. We then calculated the anomaly score of each cell after DCS normalization. The normalization and anomaly score calculation was repeated 10 times, each time with new randomly chosen 20 cells out of the 70 available cells. Receiver operating characteristic (ROC) curves were calculated for each of the 10 realizations with the cell anomaly rank as threshold. The average ROC curve is shown in in Fig. 6, curve (a). The AUC is 0.932, indicating that the anomaly score can efficiently identify cells of a phenotype different than the majority of the other cells in the dataset. As a control, we replaced the 70 endothelial cells with 928 T cells from a bone marrow sample (from SRS3805245, GSM3396161) sequenced using Illumina HiSeq 3000 and 10x chromium, but otherwise very similar to the endothelial cells in sequencing depth and median number of expressed genes per cell. As before, we chose 20 out of the 928 T cells and combined them with 1637 blood cells. The ROC curve averaged over 10 random realizations average is shown in Fig. 6, curve (b). The AUC is 0.569, indicating that cells of similar phenotype get similar score despite being sequenced in different experiments.

**Fig. 6.**
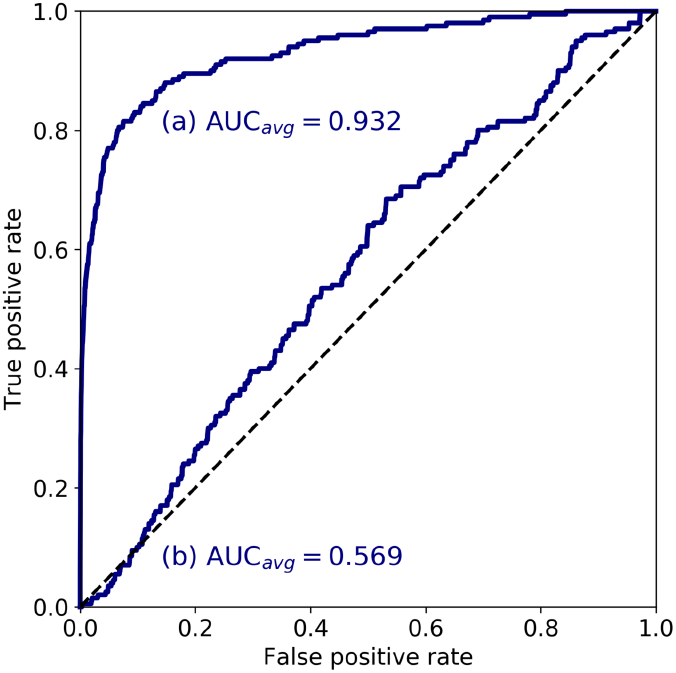
ROC curves based on the anomaly score rank for 20 cells in a set 1657 cells. The curve (a) is the average ROC for 20 endothelial cells in a majority of PBMC cells, while curve (b) is the average ROC for 20 T cells in a majority of PBMC cells.

## 4 Summary

We presented DCS, a Python-based software package for scRNA-seq data that introduces new methodologies for cell type identification, anomaly quantification, and visualization. The software provides user-friendly analysis tools for single-cell transcriptomics, starting with pre-processing and quality control, and ending with cell type annotation and downstream analysis of the annotated data. The toolkit integrates novel methodologies: (1) a tool to quantify the anomaly of cells based on isolation forest, an unsupervised machine learning algorithm for anomaly detection. We use this algorithm to detect cells that are different than the majority of other cells in a dataset, but that do not cluster together. (2) A new method for cell type identification based on a Hopfield network classifier. This method represents different cell types as attractors of the signaling dynamics in a gene network. A measured cell’s expression evolves according to this model, and its label is assigned by the attractor is converging into. (3) An enhanced version of our method for the automatic annotation of hematological cell types (pDCS). This new version uses a normalization method that accounts for markers that are known to be always unexpressed in certain cell types. Also, noise in cell type assignment has been significantly reduced in the new version. We have shown how to combine different algorithms in our toolkit to obtain a consensus score for the cell annotation. This consensus method can correctly identify cell types in synthetic mixtures of cell types. The toolkit includes extensive visualization tools, including a tool for visualizing Hopfield’s landscapes, in which cell types are represented as minima of an energy-like function representing the cellular phenotypic landscape.

## Acknowledgements

We thank Dr. Yuanfang Cai from Department of Computer Science at Drexel University for evaluating architecture and design of Digital Cell Sorter.

## Funding

This work was supported by National Institutes of Health, Grant No. R01GM122085.

## Conflict of Interest

CP and GP owns shares of Salgomed, Inc.

## Supplementary Materials

### 1.1 Q transformation on Hopfield landscapes

While Fig.2 represents a landscape obtained without the matrix *Q_μv_*, we show in Supplementary Fig. 1 the Hopfield landscape obtained using the the matrix *Q_μν_* and the *J_ij_* exactly as in Eq. 2. Note that, in this case, the Principal Component Analysis (PCA) components for the visualization has to be calculated on the transformed matrix *Q*^−1^ *ξ^T^*, where *ξ* is the matrix of genes and attractors, instead of *ξ^T^* as used in calculation of Fig. 2. From Supplementary Fig. 1 it is clear how the use of the matrix *Q_ij_* improves the separation between the attractor states, as compared to Fig. 2. The landscape is only using the first two principal components. Supplementary Fig. 2 provides a complete representation of the overlaps between the cell attractors on the top non-trivial (non-zero) principal components obtained after the *Q* transformation.

**Fig. 1.**
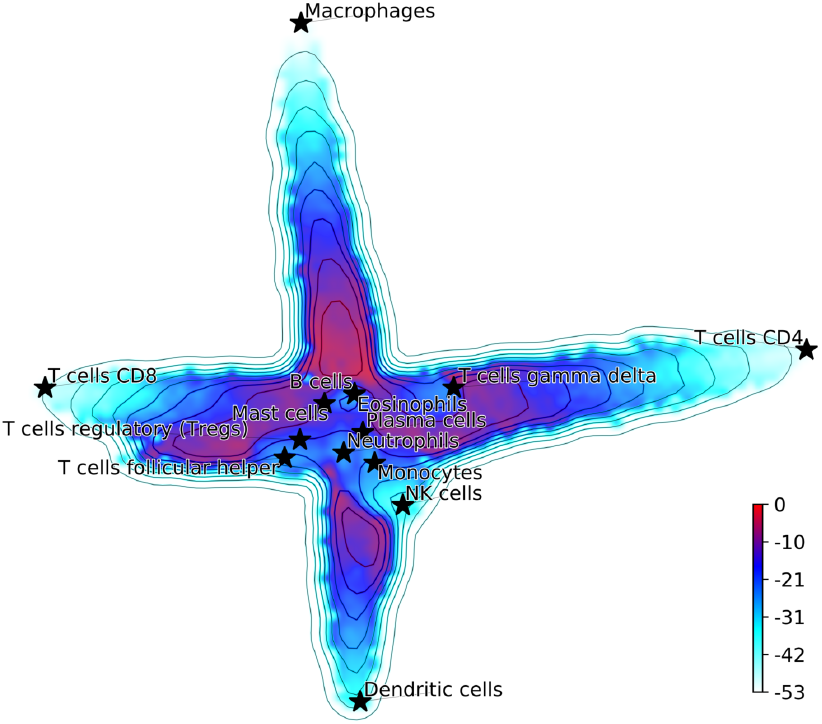
(Supplementary) Hopfield attractor landscape visualization. The points are colored according to their Hopfield energy. The *Q* transformation has improved the separation between the attractor states compared to Fig. 2.

**Fig. 2.**
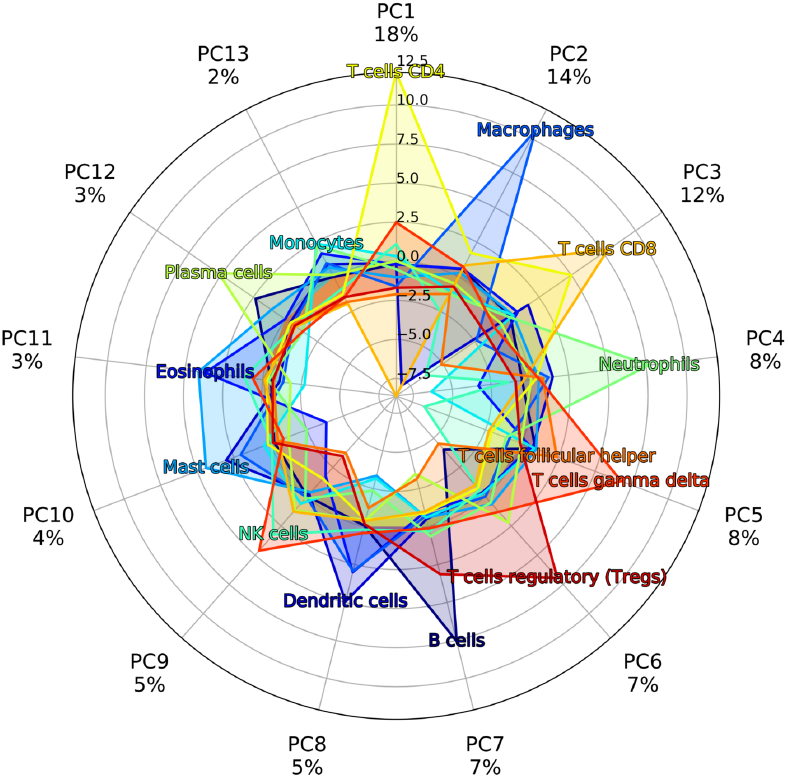
(Supplementary) Plot of the projection of different cell attractors on PCA components obtained after the *Q* transformation.

**Fig. 3.**
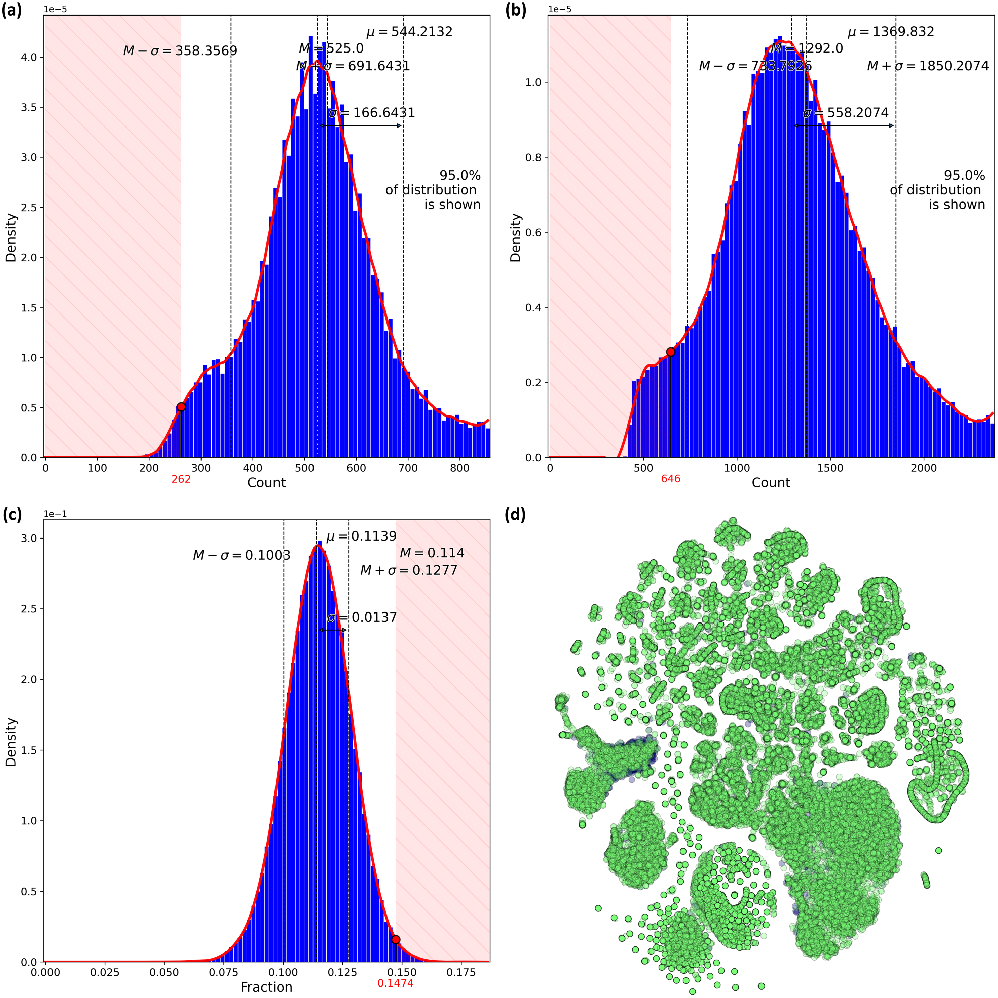
(Supplementary) Quality Control. (a) Histogram of the number of unique genes in each cell. The red line is the spline fit of the distribution. The cutoff is determined as 50% of the median of the distribution. Cells that have gene counts in the red shaded area, i.e. below the cutoff, are discarded before clustering; (b) Same as in (a) for the number of total reads per cell; (c) Histogram of fraction of mitochondrial genes in each cell. Values in the red shaded area are discarded; (d) t-SNE layout of the analyzed data where the dark blue points are cells discarded as not passing quality criteria in (a)-(c).

### 1.2 Integration with existing algorithms

#### 1.2.1 Batch correction

Data often contain noise associated to batch effects due to systematic experimental errors, collection procedures, and data handling ((4)). It is important to remove batch variations and technical noise in the data, yet preserving biologically relevant information. DCS includes a batch effect correction method, COMBAT, that has been developed specifically for microarray data, ((3, 10)) but is also suitable for single cell transcriptomics data.

Handling of missing values is particularly important when dealing with batch effects correction in scRNA-seq. In our implementation of COMBAT we record the locations of missing values, i.e. zeros in the dataset, and perform the COMBAT transformation. Then, we replace with zeros any values that were missing and became nonzero after the transformation. We also replace with zeros any values that became negative.

Supplementary Fig. 4 demonstrates how scRNA-seq data of plasma cells from the bone marrow aspirates of 12 patients (MM01-MM12) split into clusters corresponding to different batches. Processing the data with COMBAT and applying our procedure for missing values properly aligns the multi-dimensional data.

#### 1.2.2 Clustering

DCS contains functions that implement different clustering methods:

1. Hierarchical clustering. This is in general a good choice for clustering single cell datasets ((5)). However, this method becomes unfeasible when the size of the datasets goes beyond several tens of thousands cells. In this case DCS can use approximate methods such as k-means.
2. Network-based clustering methods. First a network of cells
constructed. Typically, it is a knn-graph (k nearest neighbors graph) with a cutoff k on the number of the nearest neighbors of each node. Then clusters are found using networkbased algorithms such as modularity-based community detection ((9)).
3. Spectral co-clustering ((2)). Since this method is computationally demanding, it is recommended only for small subsets of the data. We have found that spectral co-clustering can accurately fine-separate cell sub-types when used with our cell type identification algorithms. In this case, we perform first a coarse clustering using one of the two methods above (e.g. to identify all T cells in the dataset) and then we use co-clustering to obtain cell subtypes (e.g. T CD8+, T CD4+, T memory, T naive, etc.).

#### 1.2.3 Projection of high-dimensional data on 2D layout

Visualization of cell clusters can be done in DCS using different state-of-the-art methods to represent the results in a 2D layout:

1. t-SNE, Supplementary Fig. 5 (a), a well-established nonlinear method that preserves local data structure ((6)).
2. PCA, based on the first two principal components (2nd PC vs. 1st PC) of the data, see Supplementary Fig. 5 (b). This is a simple linear method that preserves the global structure of the data.
3. UMAP. This approach ((7)) has recently gained popularity because it preserves inter-cell distance in the dimensionality reduction procedure. This algorithm maintains the global structure and the continuity of the expression data. UMAP has been found to resolve cell populations and to produce equally meaningful representations compared with t-SNE ((1)). Layouts produced with UMAP are more reproducible than other methods, notably more so than those from t-SNE. An example of visualization using UMAP on PBMC dataset is shown in Supplementary Fig. 5 (c).
4. PHATE, showed in Supplementary Fig. 5 (d). This recently developed method captures local and global structure using an information-geometric distance between data points ((8)). PHATE has been found to reveal biological insights into cell developmental branches, including identification of previously undescribed subpopulations ((8)).

### 1.3 Input gene expression data

The input gene expression data is expected in one of the following formats:

- Spreadsheet of comma-separated values (csv) containing a condensed matrix in a form (“cell”, “gene”, “expr”). If there are batches in the data, the matrix has to be of the form is (”batch”, ”cell”, ”gene”, ”expr“). Column order can be arbitrary.
- Spreadsheet of comma-separated values csv where rows are genes, columns are cells with gene expression counts. If there are batches in the data the spreadsheet the first row should be ”batch“ and the second “cell”.
- Pandas DataFrame where axis 0 is genes and axis 1 are cells. If there are batched in the data, then the index of axis 1 should have two levels, e.g. (”batch”, ”cell“), with the first level indicating patient, batch or experiment where that cell was sequenced, and the second level containing cell barcodes.
- Pandas Series where the index should have two levels, e.g. (”cell”, ”gene“). If there are batched in the data the first level should be indicating patient, batch or experiment where that cell was sequenced, the second level cell barcodes, and the third level gene names.

### 1.4 Miscellaneous analysis tools

The optimized performance of our modular DCS software allows for an efficient processing of large single cell datasets. The documentation of our software is built with Sphinx at ReadTheDocs.org. Any changes to source code and python code docstrings are automatically automatically reflected at ReadTheDocs.org, and a new version of the documentation is built.

In DCS we have implemented numerous querying functions for an efficient extraction of cells based on a specific cluster or cell type. This functionality allows for an easy selection of data subsets for further analysis.

A specialized function in DCS provides across clusters comparison via a two-tailed t-test plot of individual genes. See the example of CD4 expression in PBMC data in the Supplementary Fig. 6.

Finally, DCS includes a function representing in a pie chart the role of different markers in a direct comparison between two cell types. Supplementary Fig. 7 shows an example of output for T cells versus NK cells markers. This function can be useful, for instance, to make the final decision on a cluster for which the cell type assignment has provided two possible cell types.

**Fig. 4.**
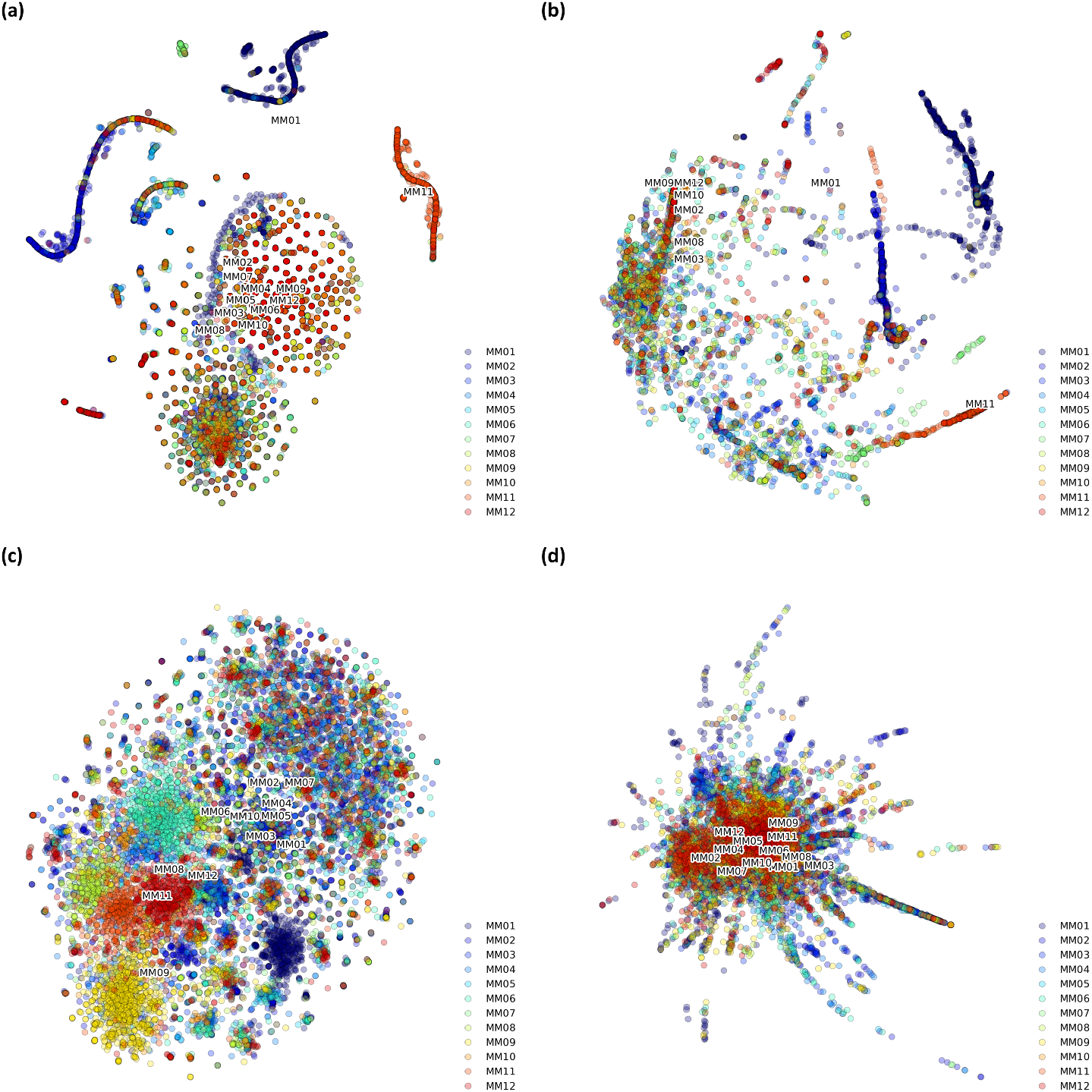
(Supplementary) scRNA-seq data of Bone Marrow plasma cells from 12 patients (MM01-MM12) split into clusters clearly showing batch-effects in (a) t-SNE layout and (b) PHATE layout. Processing these data sets with COMBAT aligns the multi-dimensional transcriptomics data mitigating batch effects as seen in (c) t-SNE layout and (d) PHATE layout.

**Fig. 5.**
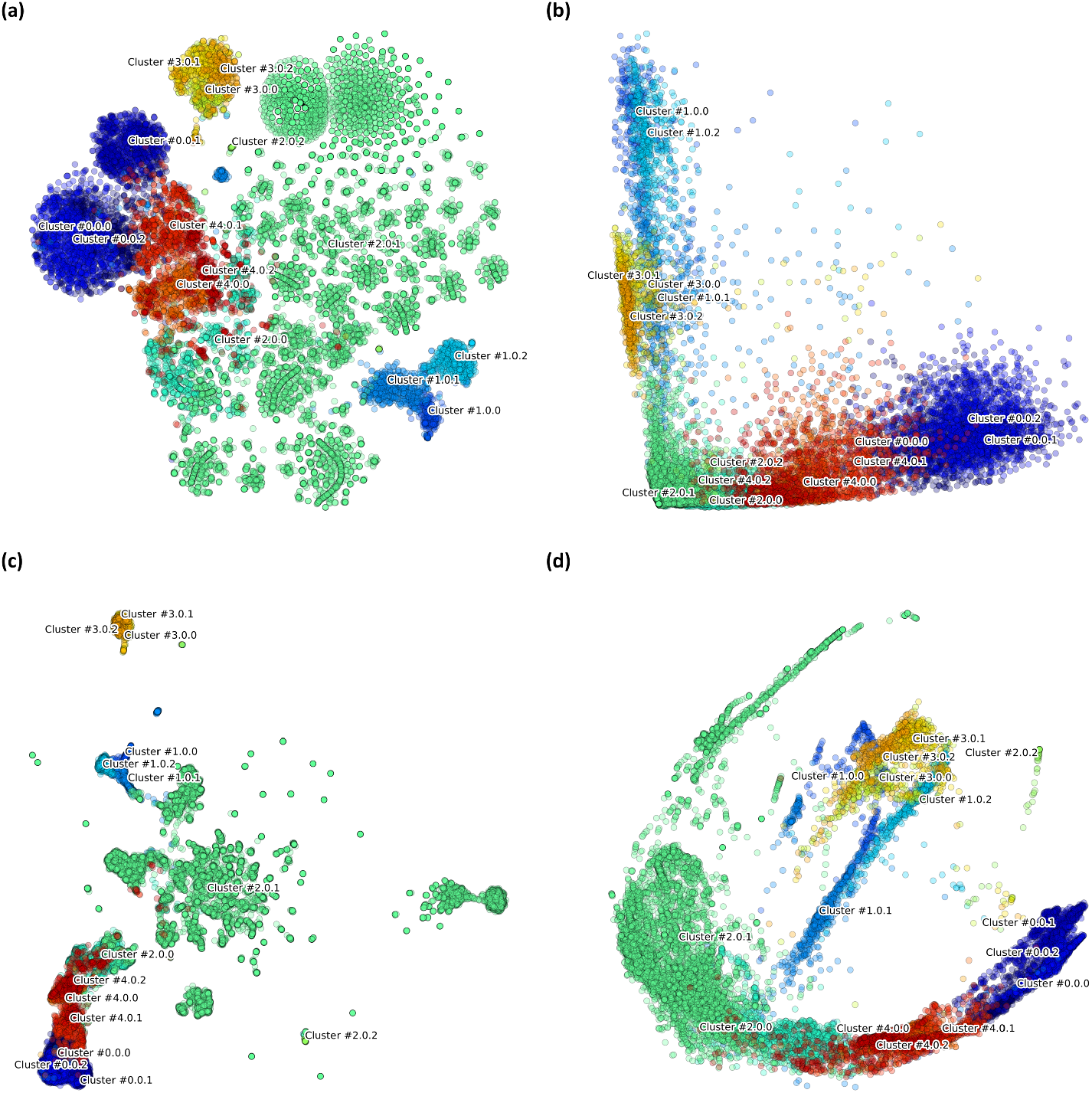
(Supplementary) Two-dimensional projection of PBMC scRNA-seq data on (a) t-SNE, (b) two largest principal components of PCA, (c) PHATE layout, and (d) UMAP layout.

**Fig. 6.**
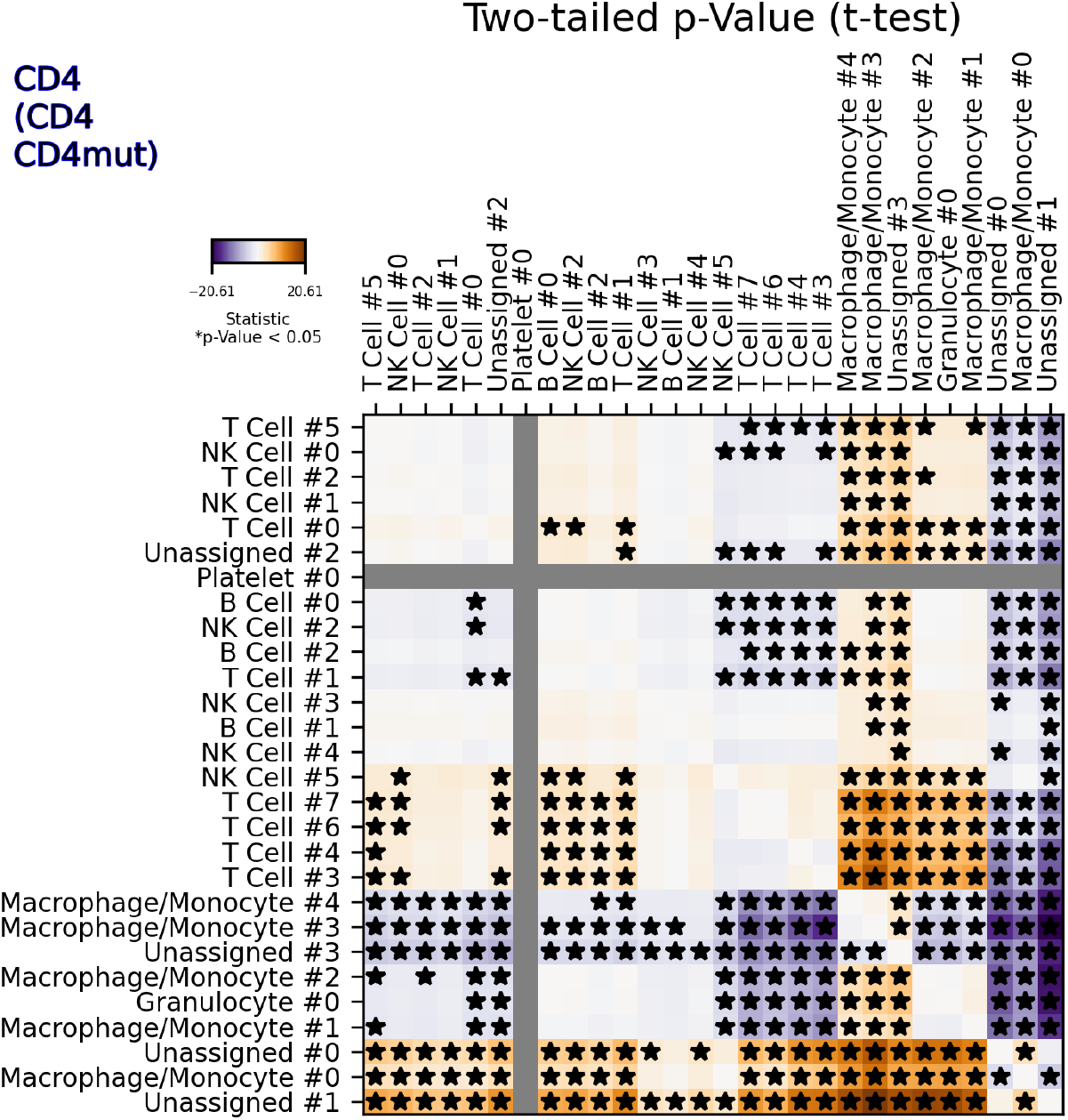
(Supplementary) Example of two-tailed t-test analysis for the CD4 gene expression in the 68k PBMC dataset. Black stars denote where this gene is significantly expressed in a two-cluster cross-comparison.

**Fig. 7.**
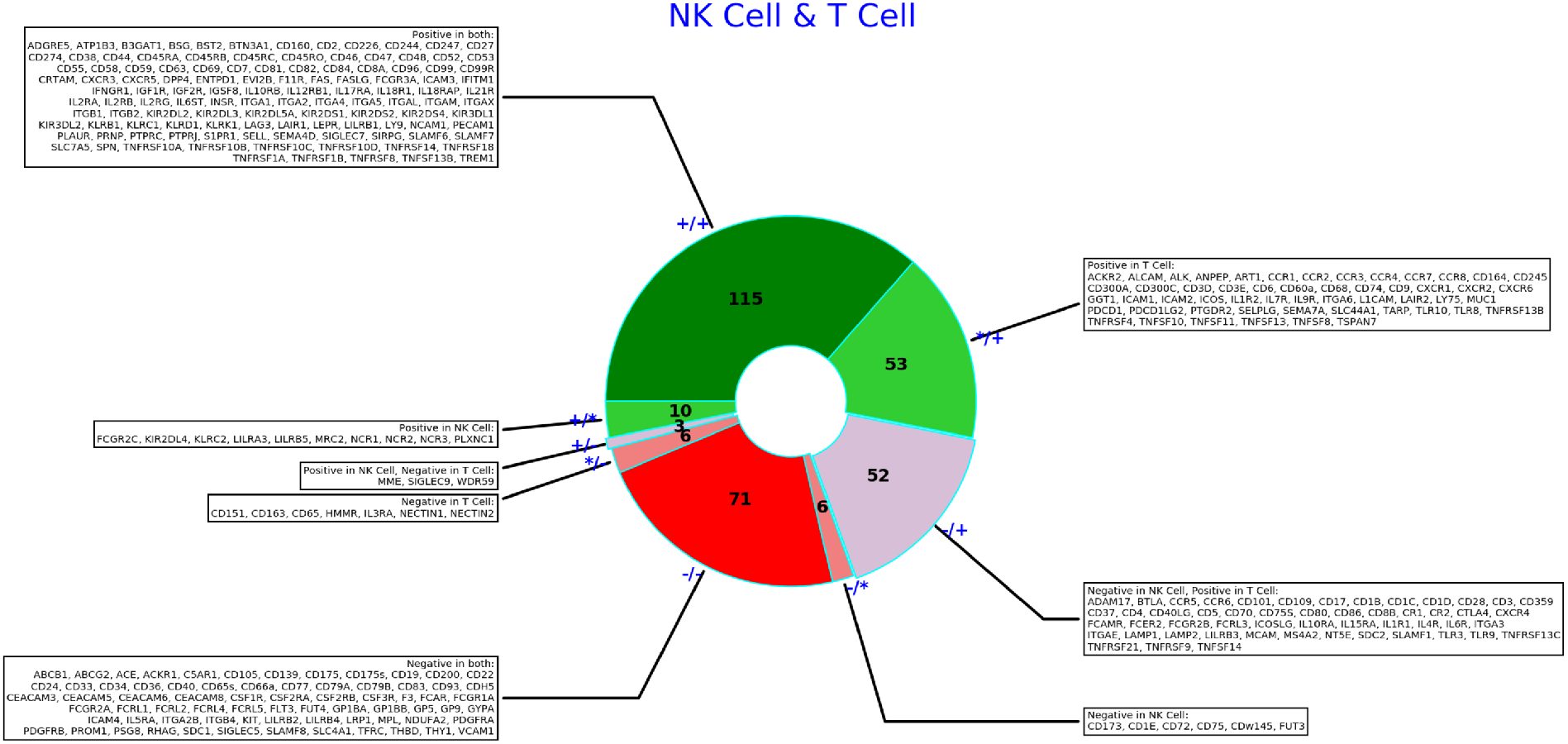
(Supplementary) Summary of T cells and NK cells markers from the “CD Marker Handbook”. The dark green sector shows markers expected to be found in both cell types, the light green sector contains markers expressed in one of the cell types. The dark red sector is for markers that are not expected to be significantly expressed in either cell types, and light red is for markers that should not be significantly expressed in one of the two cell types. Grey-shaded sector contain markers that are expected to be expressed in one of the cell type and non-expressed in the other. The number in the center is the total number of known markers for these two cell types.

**Table 1.** (Supplementary) Metadata of samples used in validation of anomaly detection.

|  |  |  |  |
| --- | --- | --- | --- |
| *Study accession* | GSM3169075 | GSM3402081 | GSM3396161 |
| *Sample accession* | SRA713577, SRS3363004 | SRA784242, SRS3822686 | SRA779509, SRS3805245 |
| *Sequencing protocol* | 10x chromium | 10x chromium | 10x chromium |
| *Sequencing instrument* | Illumina HiSeq 2500 | Illumina HiSeq 2500 | Illumina HiSeq 3000 |
| *Tissue origin of the sample* | Peripheral blood mononuclear cells | Testicular cells | Bone marrow |
| *Number of sequenced genes* | 27483 | 28403 | 30594 |
| *Number of sequenced cells* | 6008 | 6361 | 8307 |
| *Number of QC-passed cells* | 3167 | 1501 | 5928 |
| *Median of genes per cell in QC-passed cells* | 1178 | 1100 | 1106 |
| *Used cell clusters* | Cluster #0: 666 T cells,<br>Cluster #1: 564 T cells,<br>Cluster #2: 407 Monocytes | Cluster #6: 70 Endothelial cells | Cluster #0: 928 T cells |

## Notes

https://github.com/sdomanskyi/DigitalCellSorter

